# Molecular determinants of the *Bacillus subtilis* chromosome origin basal unwinding system

**DOI:** 10.1101/2022.03.18.484894

**Authors:** Simone Pelliciari, Daniel R. Burnham, George Merces, Hasan Yardimci, Heath Murray

## Abstract

Genome duplication is essential for cell proliferation and DNA synthesis is generally initiated by dedicated replication proteins at specific loci termed origins. During DNA replication initiation in bacteria, the ubiquitous DnaA protein engages both double-strand DNA (dsDNA) and single-stranded DNA (ssDNA) at the chromosome origin (*oriC*) to promote DNA duplex unwinding. While the molecular basis for DnaA binding to a specific dsDNA element (“DnaA-box”) has been established, the mechanism for DnaA binding to a specific ssDNA motif (“DnaA-trio”) is unclear. Here we define specific steps of DnaA-trio engagement by *Bacillus subtilis* DnaA. Single-molecule total internal reflection fluorescence microscopy indicates that DnaA proteins are loaded onto DnaA-trios using DnaA-boxes located on a shared DNA polymer. Chemical modification of either the phosphodiester backbone or the nucleobases revealed that three DnaA-trio repeats proximal to DnaA-boxes are necessary and sufficient to promote DnaA-dependent strand separation, and that the amino group from the central nucleobase of the DnaA-trio is critical for this reaction. Finally, based on electrophoretic mobility shift assays, we propose that during replication initiation DnaA progresses from DnaA-boxes to nucleobase recognition at DnaA-trios before engaging the phosphodiester backbone and destabilizing the DNA duplex. These results provide new molecular insight into DnaA-dependent *B*acterial *U*nwinding *S*ystem (BUS) activity at a bacterial chromosome origin.

## INTRODUCTION

Accurate transmission of genetic material is a fundamental requirement for life. In most cells, DNA replication must commence once (and only once) per division cycle to ensure rigorous coordination of genome duplication and segregation. Dysfunction of DNA replication initiation can lead to improper chromosome inheritance, disease and cell death.

Throughout the domains of life, conserved proteins containing AAA+ (*A*TPase *A*ssociated with various cellular *A*ctivities) domains assemble into dynamic multimeric complexes on DNA and direct loading of the replicative helicase (Bleichert et al., 2017). Subsequent helicase activation promotes assembly of the replication machinery that catalyses DNA synthesis. In bacteria, the toroid helicases that drive bidirectional replication from a chromosome origin are loaded around ssDNA. Bacterial chromosome origin unwinding to permit helicase loading proceeds through a mechanism involving the ubiquitous master initiation protein DnaA.

DnaA is a multifunctional enzyme composed of four distinct domains that act in concert during DNA replication initiation (Fig. S1) (Messer W., 1999). Domain IV contains a helix-turn-helix dsDNA binding motif that specifically recognizes the 9 base-pair DnaA-box (consensus 5′-TTATCCACA-3′) (Fujikawa et al., 2003; Fuller R. S., 1984; Roth A., 1995). Domain III is composed of the AAA+ motif that can assemble into an ATP-dependent right-handed helical oligomer (Erzberger et al., 2006; Erzberger et al., 2002; Schaper and Messer, 1997). Domain III also contains a ssDNA binding motif that specifically recognizes the trinucleotide DnaA-trio (consensus 3′-GAT-5′) (Duderstadt et al., 2011; Ozaki et al., 2008; Richardson et al., 2016). The DnaA^ATP^ oligomer is thought to bind the phosphodiester backbone of one DNA strand and promote destabilization of the DNA duplex by stretching the ssDNA substrate (Cheng et al., 2015; Duderstadt et al., 2011; Richardson et al., 2019). Domain II tethers domains III-IV to domain I, and domain I acts as an interaction hub that facilitates loading of the replicative helicase (Sutton et al., 1998).

Two models have been proposed for how DnaA uses its dsDNA and ssDNA binding activities to promote *oriC* unwinding (Fig. 1B). Based mainly on studies using *Escherichia coli*, in one model DnaA forms an oligomer on dsDNA that simultaneously binds ssDNA. Here a DnaA^ATP^ oligomer assembles at DnaA-boxes upstream of the *oriC* unwinding site and then interacts with a single DNA strand from downstream that is delivered by a loop mediated by the architectural protein IHF (Noguchi et al., 2015; Ozaki and Katayama, 2012; Sakiyama et al., 2017; Shimizu et al., 2016). Alternatively, based mainly on studies of *Aquifex aeolicus*, an opposing model posits that DnaA binding to dsDNA and ssDNA is mutually exclusive. Here DnaA binding to DnaA-boxes acts as a platform for directing a distinct DnaA^ATP^ oligomer onto a single DNA strand located adjacent (Duderstadt *et al.*, 2011; Duderstadt et al., 2010; Erzberger *et al.*, 2006).

**Figure 1.**
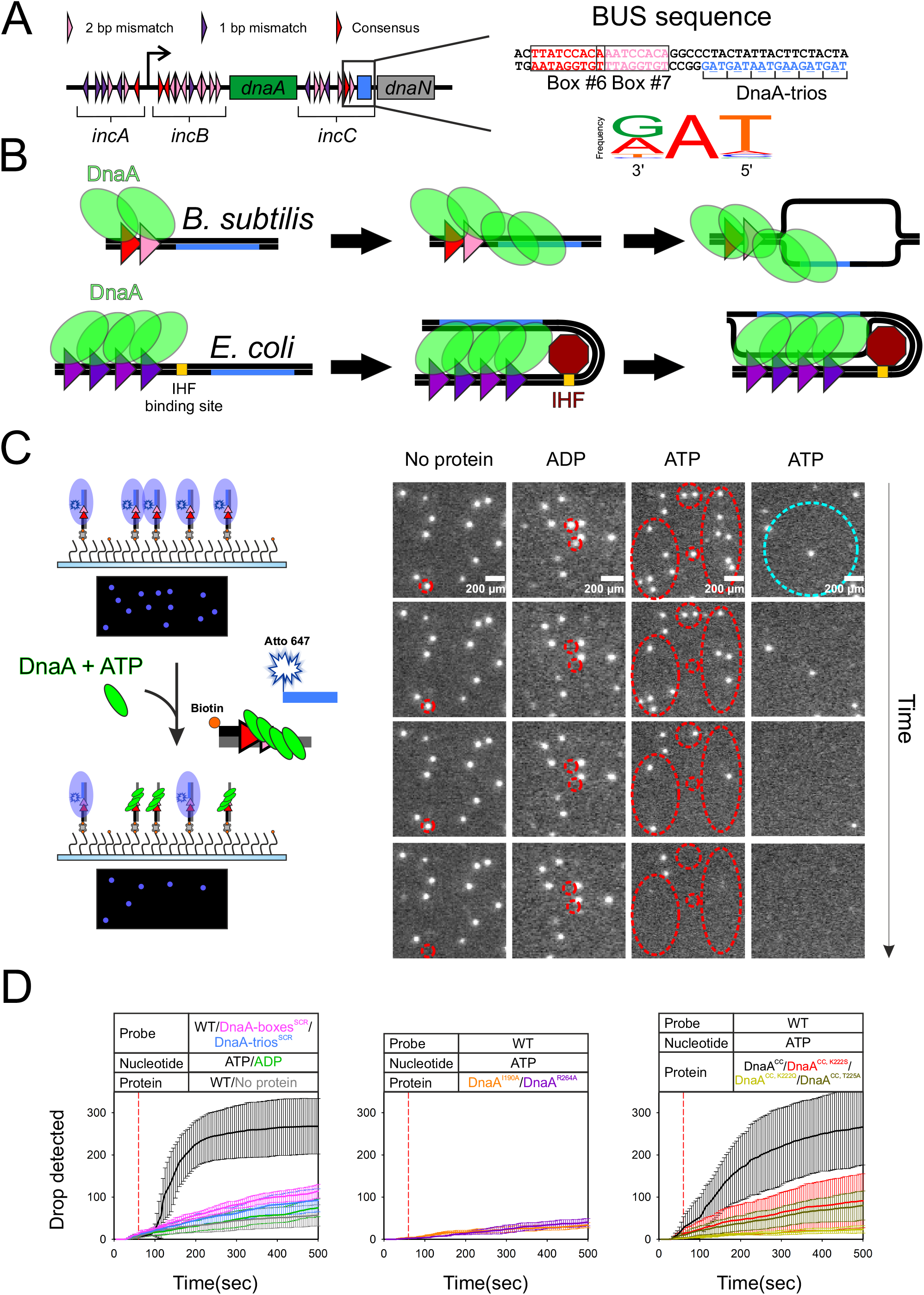
Single molecule visualization of strand separation events. **(A)** Schematic representation of the *B. subtilis* replication origin and BUS sequence, encompassing DnaA-box#6-7 and the DnaA-trios. **(B)** Proposed models of DnaA strand separation activity. **(C)** On the left, setup of single molecule microscopy experiments. On the right, fields of view from experiments performed using different conditions (indicated on top) as a function of time. Red circles indicate clusters or single spots that disappear from the field of view during the experiments. The blue circle highlights a well isolated DNA probe (radius 0.5 mm) that is not in contact with any other DNA molecules. **(D)** Graphs reporting the additive number of spots with decreased intensity over time (different experimental conditions indicated along top with colour code).

Bacterial replication origins encode information that promotes specific unwinding of the DNA duplex by DnaA (Bramhill D., 1988; Wolanski et al., 2014). Typically, bacterial origins contain multiple DnaA-boxes, flanked by the DNA unwinding site which often contains DnaA-trios and an intrinsically unstable AT-rich element (Fuller R. S., 1984; Kowalski and Eddy, 1989; Pelliciari et al., 2021; Richardson *et al.*, 2016). Comparisons of *oriC* regions from throughout the bacterial domain suggested that they are highly diverse, containing a variable number and distribution of DnaA-boxes (Luo et al., 2018; Mackiewicz et al., 2004). Historically, this heterogeneity has precluded recognition of the core chromosome origin sequence features.

Recently, a broadly conserved chromosome origin *B*asal *U*nwinding *S*ystem (BUS) was reported (Fig. 1A) (Pelliciari *et al.*, 2021). In this reaction, DnaA binding to DnaA-boxes (dsDNA binding) stimulates the assembly of DnaA^ATP^ into an oligomer that engages DnaA-trios (ssDNA binding) to promote DNA strand separation. However, it was unclear from previous studies of the BUS whether DnaA bound DnaA-boxes and DnaA-trios simultaneously or whether these interactions were separate (i.e. could not distinguish between models in Fig. 1B). Moreover, it is not currently understood how the DnaA-trios are specifically recognized by DnaA, nor is it known whether DnaA-trio binding by DnaA precedes or follows engagement of the phosphodiester backbone.

To investigate these fundamental questions, we directly observed isolated DNA strand separation reactions using single-molecule total internal reflection fluorescence (TIRF) microscopy. These experiments support a model for a DnaA^ATP^ oligomer delivered onto DnaA-trios from the adjacent DnaA-boxes. We go on to investigate the DNA determinants required for BUS function. The results are consistent with DNA backbone engagement by DnaA^ATP^ requiring prior base recognition of the DnaA-trios. Taken together, we propose a stepwise model for DNA strand separation at an *oriC* BUS.

## RESULTS

### The *B. subtilis* BUS does not require DnaA to bind dsDNA and ssDNA simultaneously

BUS activity was previously reconstituted *in vitro* using purified DnaA proteins and fluorescently-labelled oligonucleotide scaffolds (Pelliciari *et al.*, 2021; Richardson *et al.*, 2019). While these experiments defined the *oriC* sequence elements and DnaA activities required for the BUS, they could not discriminate between models of DnaA binding dsDNA and ssDNA simultaneously (which in these reactions would occur *in trans* between separate DNA molecules) versus binding these elements independently (Fig. S2). Therefore, to determine whether the BUS requires simultaneous dsDNA and ssDNA binding by DnaA, we immobilized fluorescently labelled DNA scaffolds onto a glass surface and directly visualized individual strand separation reactions using TIRF microscopy. If DnaA^ATP^ is active on these spatially separated substrates, then the BUS does not require DnaA to bind dsDNA and ssDNA simultaneously.

In this assay, DnaA^ATP^ oligomer activity results in detachment of the fluorescently labelled oligonucleotide from the immobilized scaffold and diffusion away into solution where it can no longer be visualised using TIRF illumination (Fig. 1C, strand separation assay, SSA). A DNA concentration was chosen to maintain an average distance of approximatively 15.2 μm (±7.4 μm) between immobilized DNA scaffolds, thereby preventing DnaA from binding dsDNA and ssDNA *in trans* (length of the DNA substate used here is ~30 nm). A microfluidic chamber was used to wash any unbound DNA molecules following scaffold immobilization, as well as to inject factors to start the reaction.

Addition of DnaA^ATP^ to the immobilized DNA scaffolds resulted in a steady reduction in the number of fluorescent molecules detected (234 drops/330 total, 71%), whereas the use of DnaA^ADP^ did not generate a comparable decrement (75/413, 18%) (Figs. 1C-D). The number of fluorescent signals lost in the presence of DnaA^ADP^ was equal to that in the absence of protein, suggesting that this signal is mainly background (i.e. non-specific photobleaching of the fluorophore) (Fig. 1D). To ensure that DnaA^ATP^ was promoting specific strand separation rather than non-specific scaffold disassembly, two oligonucleotides of the DNA substrate were fluorescently labelled (Figs. S3A-B). This analysis showed that DnaA^ATP^ specifically separates the fluorescently labelled oligonucleotide that complements the DnaA-trios, while leaving the remainder of the DNA scaffold intact (Figs. S3A-B). Although the ATTO^565^ fluorophore photo-bleaches rapidly, limiting its usefulness, the relatively constant signal detected with the ATTO^647^ fluorophore shows that the scaffold is not prone to disassemble on its own (Fig. S3B). Finally, DnaA-dependent strand separation events were observed on DNA scaffolds separated from each other by >500 μm (Fig. 1C, light blue circle), clearly demonstrating that interactions between DNA molecules *in trans* is not required.

To determine whether the single-molecule strand separation reactions required the same *oriC* sequences and DnaA activities defined for the BUS using bulk biochemical assays (Pelliciari *et al.*, 2021; Richardson *et al.*, 2019), we tested mutated DNA scaffolds and DnaA protein variants. Scrambling the DNA sequence of either the DnaA-boxes or the DnaA-trios significantly decreased the number of observed unwinding events (Fig. 1D, DnaA-boxes^SCR^ 92/609, 15%, DnaA-trios^SCR^ 114/706, 16%), showing that the DnaA-dependent strand separation reaction was specific. DnaA variants that lack key residues for either protein oligomerization (DnaA^R264A^; the “arginine finger”) or ssDNA binding (DnaA^I190A^, DnaA^K222S^, DnaA^K222Q,^ DnaA^T225A^) were unable to promote strand separation of BUS scaffolds (Fig. 1D, DnaA^R264A^ 40/353, 11%, DnaA^I190A^ 33/309, 11%, DnaA^K222S^ 91/370, 25%, DnaA^K222Q^ 23/144, 16%, DnaA^T225A^ 80/451, 18%), consistent with the proposed mechanism for ATP-dependent DNA stretching by DnaA (Duderstadt *et al.*, 2011). We confirmed that the DnaA^K222S^, DnaA^K222Q,^ and DnaA^T225A^ variants retained the ability to bind ATP (Fig. S4A) and assemble into an ATP-dependent oligomer in solution (Fig. S4B) (assessed previously for DnaA^I190A^ and DnaA^R264A^ (Pelliciari *et al.*, 2021)). Note that all assays performed using DnaA^K222S^, DnaA^K222Q,^ DnaA^T225A^ utilized a version of DnaA (DnaA^CC^) carrying a pair of cysteine residues within the AAA+ motif to facilitate bismaleimidoethane crosslinking (Fig. 1D, 290/445, 65%) (Scholefield et al., 2012). Finally, we tested a DnaA variant lacking domain I-II (DnaA^104-446^) and observed that the rate of strand separation was similar to wild-type (Fig. S5A-B, 254/387, 66%), indicating that these domains are not required.

Taken together, single-molecule analysis of the *B. subtilis* BUS indicates that DnaA strand separation activity does not require the protein to bind dsDNA and ssDNA simultaneously. The results are consistent with a model where a DnaA^ATP^ oligomer is loaded onto DnaA-trios from the adjacent DnaA-boxes.

### Location of critical phosphodiester bonds for the BUS

The DnaA^ATP^ oligomer is proposed to engage and stretch the phosphodiester backbone of a single DNA strand to promote chromosome origin opening (Duderstadt *et al.*, 2011). To investigate the extent of this DNA backbone interaction for BUS function, DNA scaffolds were designed with phosphorothionate linkages between various nucleotides of the DnaA-trios (Fig. 2B). Here DnaA strand separation activity was assayed in bulk using scaffolds containing a *B*lack *H*ole *Q*uencher (BHQ2) and a fluorophore (Cy5); separation of the fluorescently labelled oligonucleotide from the scaffold is detected using a plate reader (Pelliciari *et al.*, 2021).

**Figure 2.**
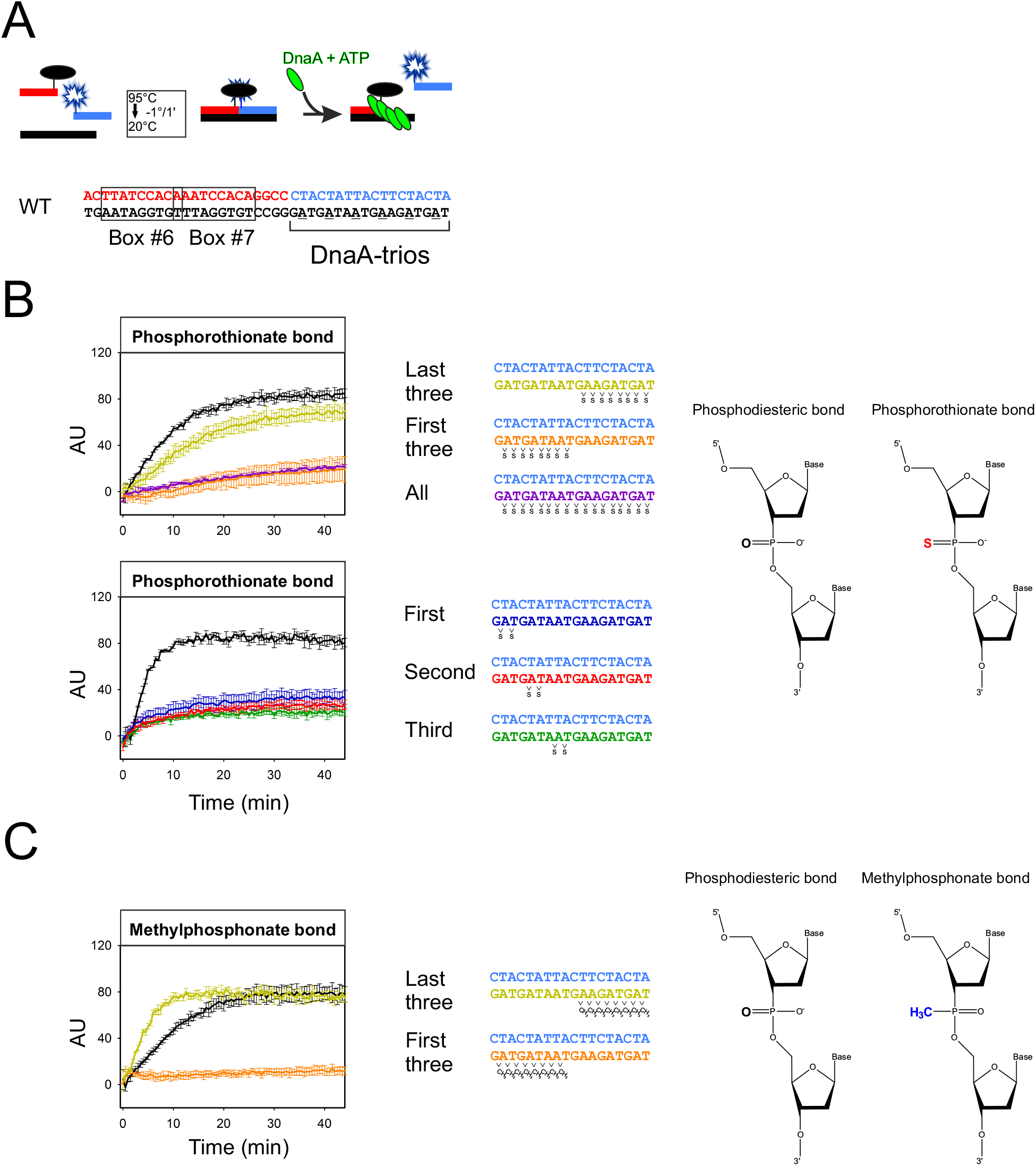
DNA backbone is fundamental for DnaA strand separation activity. **(A)** Schematic representation of BHQ strand separation assay. The wild-type BUS sequence is shown for comparison. **(B)** BHQ-SSA performed with substrates harbouring different phosphorothionate modifications within DnaA-trios. Positions of chemical modifications are indicated to the right. **(C)** Assay performed with DNA scaffolds containing methylphosphonate modification within DnaA-trios. Positions of chemical modifications are indicated to the right.

Phosphorothionate bonds throughout the DnaA-trios inhibited DnaA strand separation activity, as did phosphorothionate bonds in DnaA-trios located proximal to the DnaA-boxes (DnaA-trios#1-3) (Fig. 2A-B). In contrast, phosphorothionate bonds were well tolerated within the DnaA-trios located distal to the DnaA-boxes (DnaA-trios#4-6) (Fig. 2B). Moreover, phosphorothionate bonds throughout the DNA strand complementing the DnaA-trios did not inhibit the reaction, consistent with DnaA only needing to engage one DNA strand (Fig. S6). Focused replacements of phosphorothionate bonds within the first three DnaA-trio motifs confirmed the importance of each backbone contact to the strand separation reaction (Fig. 2B). Similar results were obtained using methyl phosphonate linkages within the DnaA-trios (Fig. 2C). Together these results suggest that the most critical interactions between DnaA^ATP^ and the phosphodiester backbone occur closest to the DnaA-boxes.

### Location of critical nucleobases for the BUS

A similar experimental approach as described above was followed to localize the most critical nucleobases within the DnaA-trios, except here trinucleotide motifs were substituted by their complementary sequence. Strand unwind assays showed that mutation of any of the three DnaA-trios proximal to the DnaA-boxes significantly inhibited the reaction (Fig. 3A-B). In contrast, mutation of either DnaA-trio#4 alone or DnaA-trio#4-6 together did not inhibit the reaction (Fig. 3B). These results suggest that the most critical interactions between DnaA^ATP^ and the DnaA-trios occur closest to the DnaA-boxes at DnaA-trios#1-3.

**Figure 3.**
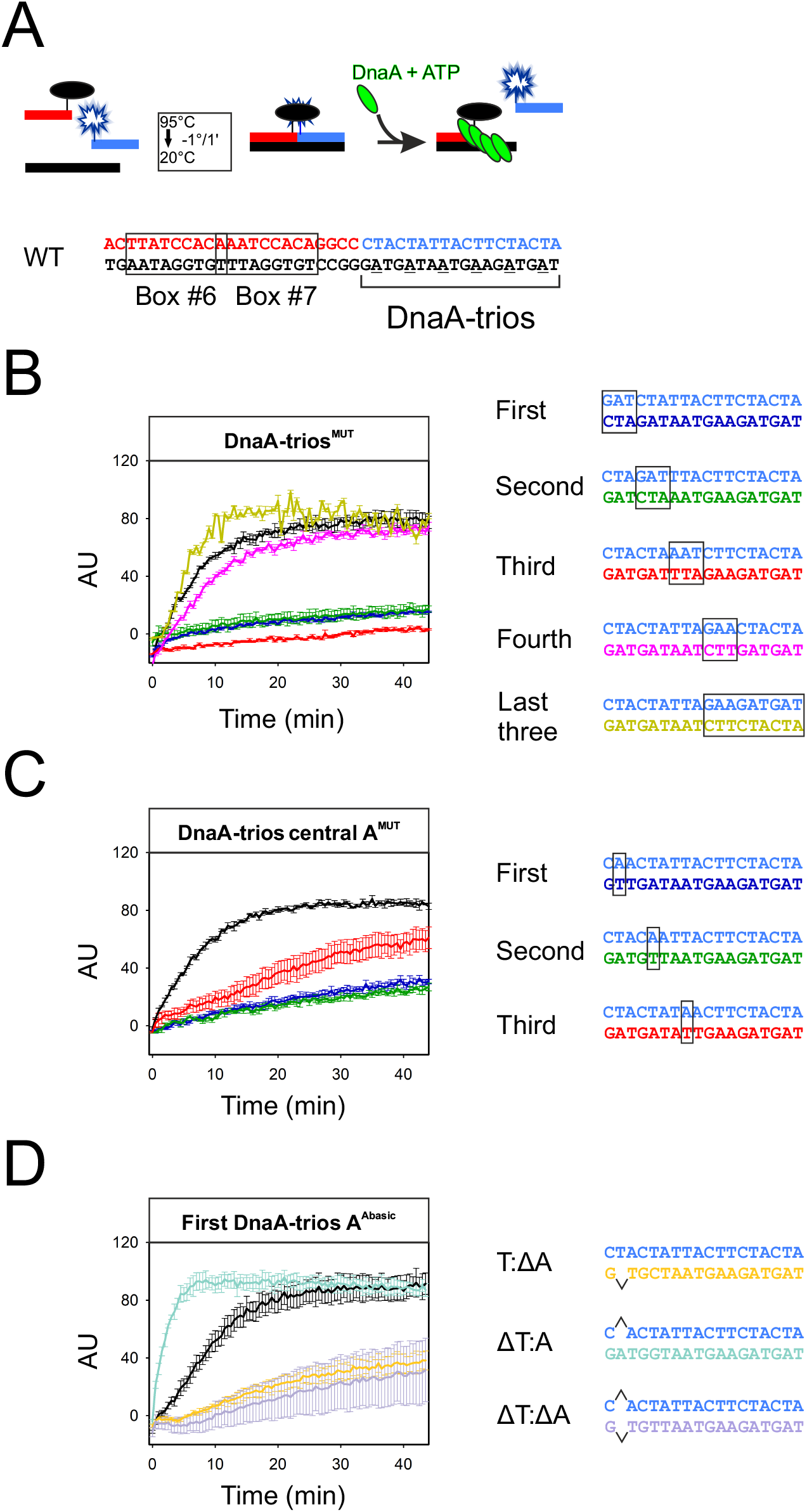
The central adenine of the first DnaA-trios is fundamental for DnaA activity. **(A)** Schematic representation of BHQ strand separation assay. The wild-type BUS sequence is shown for comparison. **(B)** BHQ-SSA performed with sequential substitution of DnaA-trios with their complementary sequence. Positions of mutations are indicated to the right. **(C)** BHQ-SSA performed substituting the central adenine of the first three DnaA-trios with the complementary nucleotide. Positions of mutations are indicated to the right. **(D)** BHQ-SSA performed using probes containing abasic sites (nucleotide substituted with THF) in the central position of the first DnaA-trio. Positions of abasic sites are indicated to the right.

### The central adenine within a DnaA-trio is required for the BUS

The DnaA-trio is a degenerate ssDNA binding motif, with the central adenine being the most highly conserved (Pelliciari *et al.*, 2021). To assess the functional importance of this nucleobase position for DnaA activity, each central A:T base pair within DnaA-trio#1-3 was flipped. All transversions significantly reduced the rate of DnaA-dependent strand separation (Fig. 3C), demonstrating that the central adenine is critical for BUS function.

To determine whether it was the absence of adenine or the presence of thymine at this position that inhibited the BUS, we constructed DNA scaffolds containing tetrahydrofuran (THF; abasic mimetic) linkages at the central position of the DnaA-trio closest to the DnaA-boxes. Removal of the adenine nucleoside significantly impaired the BUS, and this was observed whether the complementary thymine nucleoside was present or not (Fig. 3D). Notably, the reaction rate was significantly increased when the central adenine was unpaired (Fig. 3D), likely due to the lower melting temperature of the THF containing oligonucleotide (Table S1). These results indicate that the presence of adenine within the DnaA-trio closest to the DnaA-boxes is critical for the DnaA-dependent strand separation reaction.

### The central nucleobase within a DnaA-trio requires an amine functional group

The observation that the strand separation reaction readily occurred when the central adenine of DnaA-trio#1 was unpaired (Fig. 3D) allowed the opportunity to substitute this nucleobase without the complications of either mispairing with the native thymine or maintaining base pairing but introducing distinct hydrogen bonding and base stacking interactions.

The pivotal adenine was replaced by each of the natural nucleobases and strand separation assays were performed to test DnaA activity. While substitution with either guanine, thymine or uracil decreased the rate of strand separation, strikingly, substitution with the pyrimidine cytosine resulted in a wild-type rate (Fig. 4B). Both adenine and cytosine bases contain a primary amine group (linked to C_6_ for adenine, C_4_ for cytosine) that faces the major groove and acts as an electron donor. To test the hypothesis that this functional group is involved in the DnaA-dependent strand separation reaction, several modified nucleobases were tested at this central position of the DnaA-trio (adenine was replaced by hypoxanthine, substituting the C_6_ amine with a hydroxyl group; cytosine was replaced by 5-methylisocytosine, swapping the amino and carbonyl groups at positions 2 and 4; guanine was replaced by isoguanine, swapping the amino and carbonyl groups at positions 2 and 6 (Fig. 4B)). Hypoxanthine and isocytostine decreased the rate of strand separation compared to adenine and cytosine, respectively (Fig. 4B). Both of these nucleobase modifications remove the functional amine group (Fig. 4B). In stark contrast, isoguanine increased the rate of unwinding compared to guanine (Fig. 4B). In this case, the nucleobase modification adds a functional amine group. Taken together, these results support the model that the amine functional group facing the major groove plays a positive role during the DnaA-dependent strand separation reaction.

**Figure 4.**
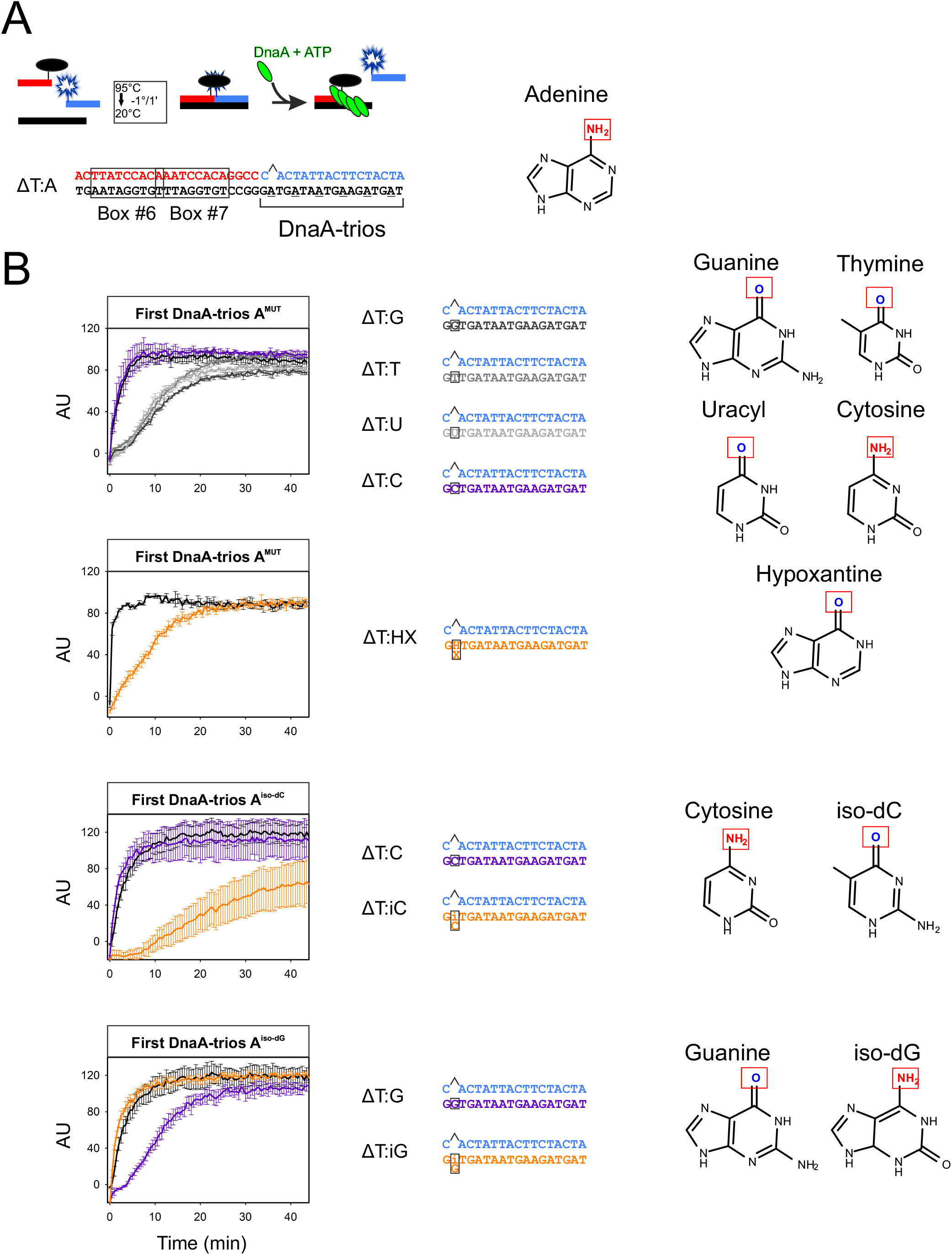
Amine group on adenine C6 is fundamental for DnaA-trios recognition. **(A)** Schematic representation of BHQ strand separation assay. The wild-type BUS sequence is shown for comparison. The critical nitrogen group of adenine is indicated. **(B)** BHQ-SSA performed with several nucleotide substitutions at the central adenine of the first DnaA-trio, in the context of an abasic site at the complementary position. Positions of mutations and structure of modified bases are indicated to the right.

### Stepwise assembly of DnaA from DnaA-trios to the phosphodiester backbone

The data collected for the BUS suggests a primary importance of DnaA-trio#1-3 proximal to the DnaA-boxes, regarding both the nucleobase base sequence of the trinucleotide motif and the composition of the backbone. We hypothesized that these two DNA interactions occur sequentially. To investigate this model, we performed electrophoretic mobility shift assays (EMSA) on multicolour DNA scaffolds containing either backbone modifications or mutated DnaA-trios. The two fluorescently labelled oligonucleotides allowed us to follow both components of the BUS scaffold and to detect DNA strand separation (Fig. 5A).

**Figure 5.**
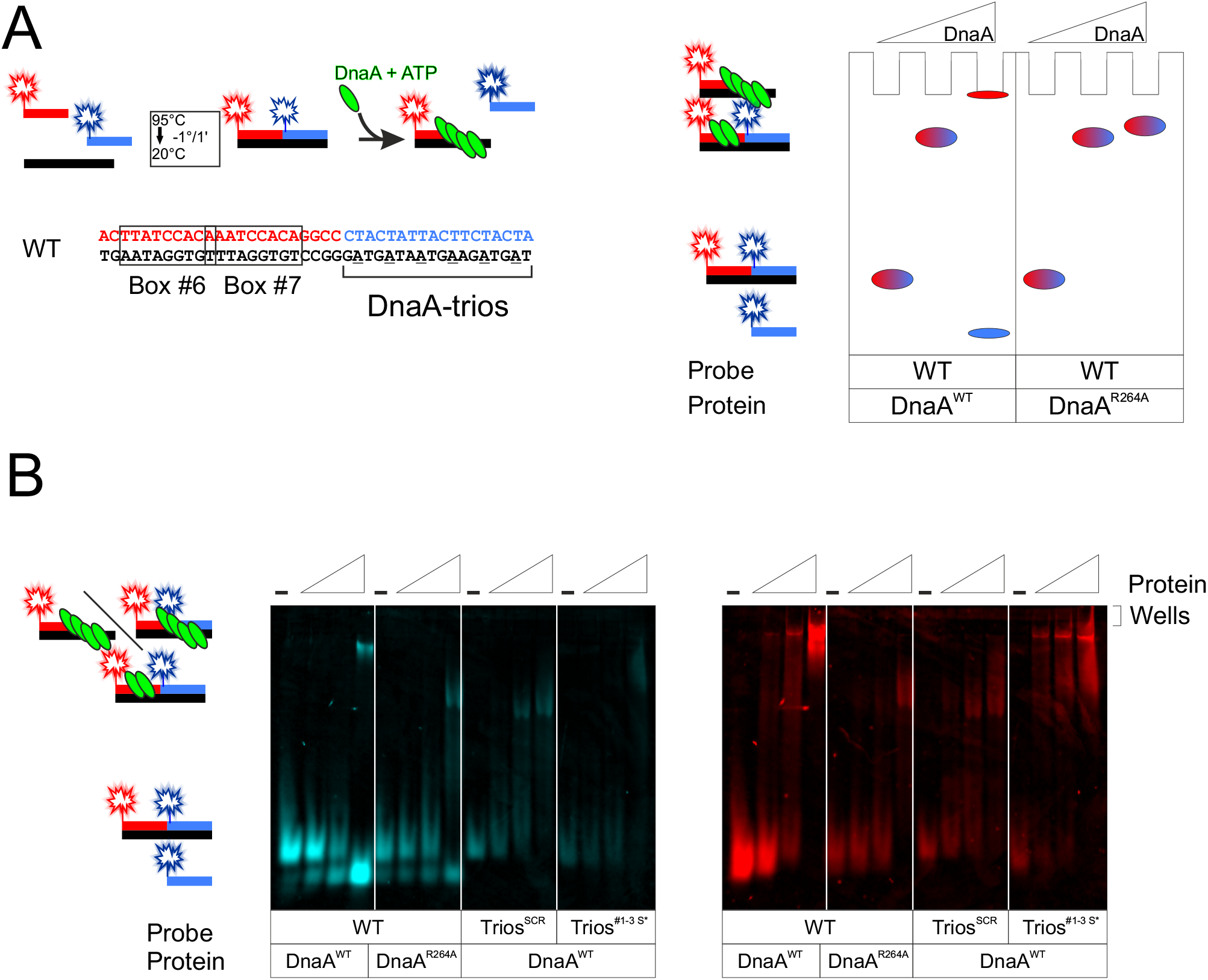
EMSA reveal the sequential order of events that leads to strand separation. **(A)** Schematic representations of the probes used and of the EMSA. **(B)** Fluorescent signals from DNA scaffolds containing two labelled oligonucleotides. The separated strand is labelled with Cy5 on the left (cyan) while the annealed strand is labelled with Cy3 on the right (red). The different high molecular weight species are indicated on the left with a schematic representation. Dash on top indicate absence of protein in the reaction and the triangle indicate the increasing concentration of DnaA used in each specific lane (250, 500, 1000 nM). Experimental conditions for each set are reported on the bottom of the figures.

Previous EMSAs showed that although DnaA^ADP^ and DnaA^ATP^ both bind to the wild-type BUS scaffold, DnaA^ADP^ generates a single shifted species that can enter the gel, while DnaA^ATP^ generates a larger nucleoprotein complex that remains near the well, in addition to liberating the fluorescently labelled strand complementing the DnaA-trios (Fig. 5B) (Richardson *et al.*, 2019). Because the single shifted species observed with DnaA^ADP^ was dependent upon DnaA-boxes but not DnaA-trios, this result implies that the DnaA^ATP^ generates a larger nucleoprotein complex that migrates poorly into the gel and is competent for strand separation (Richardson *et al.*, 2019). Consistent with this interpretation, the DnaA^R264A^ arginine finger variant that cannot assemble into an ATP-dependent oligomer produced a smaller nucleoprotein complex that enters the gel (Figs. 5A-B).

When the EMSA was performed on a DNA scaffold with scrambled DnaA-trios, the resulting DnaA^ATP^ nucleoprotein complex was observed to enter the gel (Fig. 5B). This is consistent with the notion that the DnaA^ATP^ oligomer cannot bind stably to mutated DnaA-trios, resulting in the formation of a smaller nucleoprotein complex. Alternatively, when the EMSA was performed on a DNA scaffold containing phosphorothionate modifications in the backbone, the resulting DnaA^ATP^ nucleoprotein complexes entered the gel poorly, with many appearing stuck near the well (Fig. 5B). This indicates that DnaA^ATP^ oligomer assembly does not require the backbone interaction. Taken together, the results suggest a sequential series of events during the DnaA strand separation reaction, where DnaA^ATP^ must first recognize nucleobases of the DnaA-trios before transitioning to engage and stretch the phosphodiester backbone.

## DISCUSSION

### Molecular determinants of the *B. subtilis* chromosome origin BUS

This work identifies several new molecular properties of the *B. subtilis* BUS that, together with previous works, allow us to propose a stepwise model for assembling DnaA nucleoprotein complexes at bacterial chromosome origins utilizing this strand separation mechanism.

In the BUS, DnaA^ATP^ binding to DnaA-trios requires the proximal DnaA-boxes, suggesting that DnaA is localized to *oriC* by the dsDNA binding sites before assembling into an oligomer on a single DNA strand (Pelliciari *et al.*, 2021; Richardson *et al.*, 2016; Richardson *et al.*, 2019). In the TIRF microscopy experiments reported above, these two DNA binding events must occur *in cis* on a single DNA polymer. Bulk strand separation assays indicate that DnaA requires both nucleobase specific and backbone interactions with at least three consecutive DnaA-trios for strand separation, and EMSAs suggest that nucleobase recognition precedes backbone engagement. These observations are considered along with the short length of the DNA scaffolds used characterize the BUS *in vitro*, the proximity of DnaA-boxes to DnaA-trios, the x-ray structure of a DnaA^ATP^ oligomer binding a ssDNA substrate (Duderstadt *et al.*, 2011), and the fact that domains III-IV of DnaA are sufficient for the BUS (Figs. S5B-D) strand separation.

Taken together, we propose the following model for DnaA activities within the BUS (Fig. 6). (1) DnaA_1_ binds DnaA-box#6 (consensus) using domain IV. (2) DnaA_2_ protein binds to DnaA-box#7 using domain IV. DnaA_2_ can also interact with DnaA_1_ through its AAA+ motif. Note that DnaA-box#7 is non-consensus and overlaps with DnaA-box#6; for these reasons we speculate that the DnaA_2_ domain IV interaction at DnaA-box#7 is unstable. (3) DnaA_2_ protein disengages domain IV from DnaA-box#7 and engages DnaA-trio#1 using domain III. Here DnaA-trio#1 binding would be stabilized by ATP-dependent AAA+:AAA+ interactions. Moreover, structural analysis of *Aquifex aeolicus* DnaA protein bound to ssDNA showed that domain IV of one monomer packs against the AAA+ domain of the adjacent monomer to provide an additional contact point. Interestingly, in the crystal structure this domain IV protein:protein interaction places the helix-turn-helix motif in an orientation that would preclude dsDNA binding (Duderstadt *et al.*, 2010; Erzberger *et al.*, 2006), potentially assisting in the transition of DnaA_2_ from binding DnaA-box#7 to binding DnaA-trio#1. (4) DnaA_3_ protein binds to the AAA+ domain of DnaA_2_ and DnaA-trio#2. (5) DnaA_4_ protein binds to the AAA+ domain of DnaA_3_ and DnaA-trio#3. (6) An ATP-dependent oligomer composed of DnaA_2-4_ engages the phosphodiester backbone along DnaA-trios#1-3, stretching the single strand to promote *oriC* opening.

**Figure 6.**
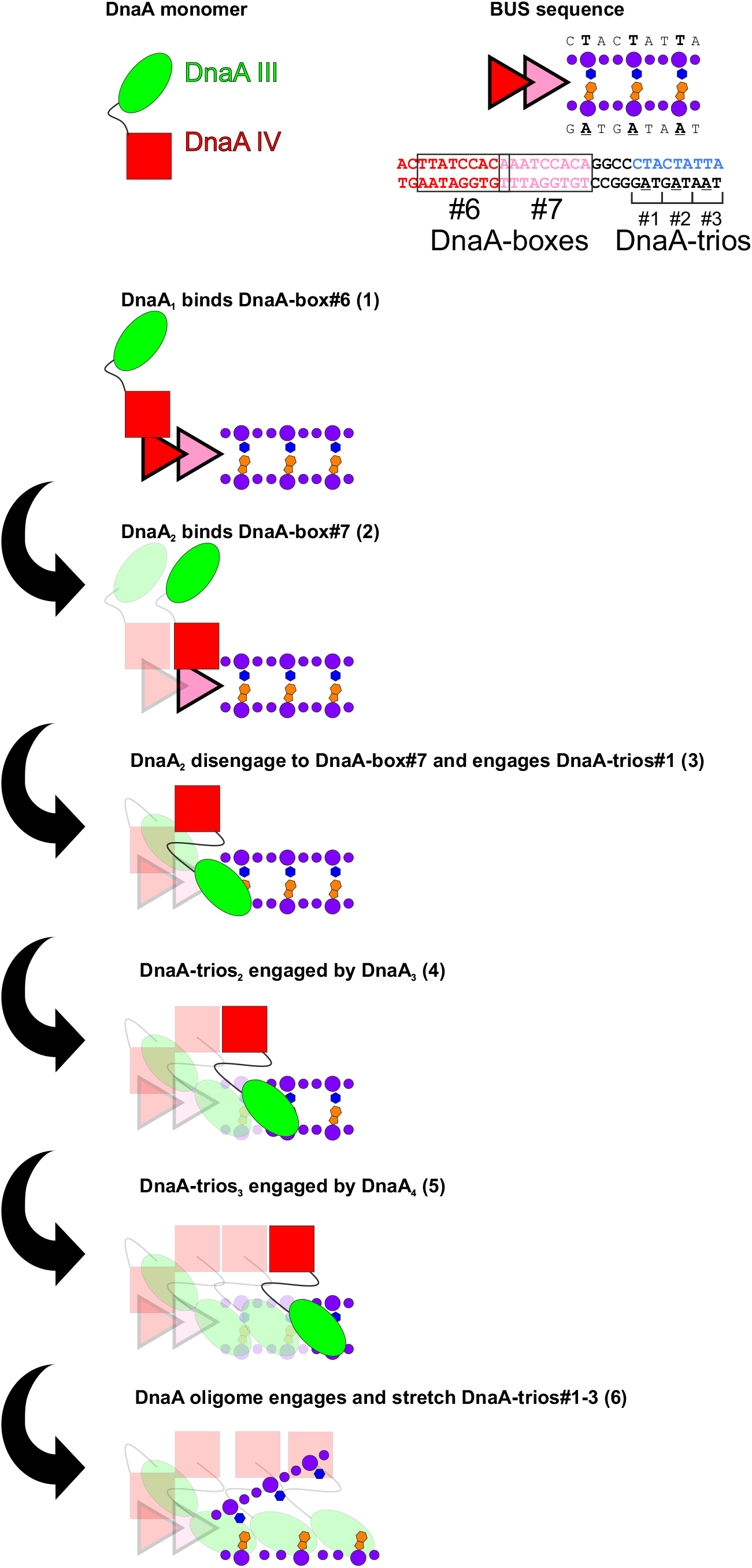
Proposed model for DnaA activities during DNA strand separation at the *B. subtilis* BUS.

Importantly, this minimal model agrees with experimental data showing that a minimum of three DnaA-trios was necessary for *B. subtilis* viability (Richardson *et al.*, 2016). Moreover, a previous bioinformatic search for BUS elements within bacterial chromosome origins set a threshold requirement of three DnaA-trios (Pelliciari *et al.*, 2021). Therefore, the model described here would apply to all putative BUS systems identified.

### “We don’t talk about supercoiling, no, no, no”

The BUS mechanism described above was primarily derived using assays that employed short oligonucleotide substrates (Duderstadt *et al.*, 2011; Pelliciari *et al.*, 2021; Richardson *et al.*, 2016; Richardson *et al.*, 2019). While this approach proved informative and the proposed model agrees well with known properties of the BUS, it is admittedly simplistic. Critically, the role of DNA supercoiling is not explored in these reactions. We suspect that local DNA topology could either alter the structure of DnaA nucleoprotein complexes or destabilize the DNA duplex to facilitate DnaA oligomer assembly and phosphodiester backbone stretching.

### Does a continuous DnaA^ATP^ oligomer bridge between DnaA-boxes and DnaA-trios?

Assays *in vitro* suggest that a DnaA^ATP^ oligomer can promote DNA strand separation in the absence of DnaA-boxes (Duderstadt *et al.*, 2011). In the BUS, DnaA-boxes are required, but their precise roles remain unclear. The DnaA-boxes would localize DnaA protein near DnaA-trios to promote specific DnaA^ATP^ oligomer assembly, but it is also possible that a continuous oligomer of DnaA^ATP^ is constructed between DnaA-boxes and DnaA-trios. DnaA_1_ bound to DnaA-box#6 could act as a brace for the DnaA^ATP^ oligomer engaging the phosphodiester backbone to push against. This could provide an additional force to drive DNA duplex opening.

### How does DnaA recognize the DnaA-trios?

It has been established that DnaA-trios are essential for DnaA strand separation activity and that they specify DnaA^ATP^ oligomer formation on ssDNA (Pelliciari *et al.*, 2021; Richardson *et al.*, 2016). However, the molecular basis for DnaA-trio recognition by DnaA was unknown, as the crystal structure of the *A. aeolicus* DnaA^ATP^ oligomer bound to ssDNA shows that each protein monomer contacts the DNA exclusively through the phosphodiester backbone (Duderstadt *et al.*, 2011). Here we elucidated that a functional nitrogen group on the central nucleobase of the DnaA-trio is pivotal for DnaA^ATP^ oligomer assembly and DNA strand separation. This is the first chemical determinant for DnaA-trio recognition by DnaA and should be considered in future molecular models for strand specific DnaA assembly.

### How is the DnaA^ATP^ oligomer orientated on DnaA-trios?

The DnaA^ATP^ oligomer displays polarity (Erzberger *et al.*, 2006). It is currently unclear which orientation the DnaA^ATP^ adopts when bound to ssDNA (Duderstadt *et al.*, 2011), but it is assumed that DnaA bound to a DnaA-box dictates this decision. One model proposes that DnaA would orientate its arginine finger residue to face away from the DnaA-boxes and towards the unwinding region, allowing an interaction with a dedicated AAA+ chaperone that guides helicase loading around ssDNA (Mott et al., 2008). A competing model proposes that DnaA would orientate its arginine finger residue to face towards the DnaA-boxes and away from the unwinding region (Noguchi *et al.*, 2015). Determining the orientation of the DnaA oligomer will have important implications for the BUS, regarding whether DnaA1 bound to DnaA-box#6 requires ATP to initiate oligomer formation, and whether the downstream interaction with the AAA+ helicase loader is plausible.

### Perspective

The functional model of the *B. subtilis* BUS provides a framework for investigating DNA replication initiation in diverse bacterial species. Moreover, the bacterial DNA replication machinery is an attractive subject for drug development, because the process is essential, the bacterial proteins are distinct from functional eukaryotic analogues, and no current antibiotics in clinical use target this pathway (Kaguni, 2018; Yi and Lu, 2019). Obtaining a clearer view of the bacterial DNA replication initiation mechanism will facilitate development of small molecule inhibitors.

## MATERIALS AND METHODS

### Media and chemicals

Nutrient agar (NA; Oxoid) was used for routine selection and maintenance of *E. coli* strains. For protein expression, cells were grown using 2X YT medium (16g Tryptone, 10g Yeast Extract, 5g NaCl for 1L) (Sambrook et al., 1989). Supplements were added as required: 1 mM IPTG, 100 μg/ml ampicillin. Unless otherwise stated all chemicals and reagents were obtained from Sigma-Aldrich.

### Glass coverslip functionalization

Glass coverslips (22 x 50 mm; VWR) were functionalized in clean staining jars and all solutions were pre-filtered. Glass surfaces were sonicated four times, 30 minutes each, in a water bath (GT SONIC-D6). After each sonication step, coverslips were washed 3 times with milliQ water. Solutions for sonication were: EtOH (100%), KOH (1 M), EtOH (100%), KOH (1 M). After the final sonication step, jars were washed five times with milliQ water, filled with KOH (1 M), and left to stand overnight. Jars were washed three times with milliQ water and then three times with acetone, prior to drying the coverslips using a N_2_ gun. A plasma cleaner was used to clean the coverslips (0.5 torr, HIGH setting for 5 min; Harrick Plasma cleaner 115 V) before placing them back in staining jars filled with acetone. To silanize the glass surfaces, coverslips were incubated in a 2% solution of (3-Aminopropyl)triethoxysilane (APTES, Sigma-Aldrich) in acetone with agitation for two minutes. The reaction was stopped by pouring water directly into the jars. Three 0.2 x 4 mm chambers were prepared using a double-sided adhesive (TESA 4965) and applied on the silanized glasses after drying them with a N_2_ gun. A 46 μM solution of PEG/biotin-PEG (in a 56:1 ratio; Laysan Bio inc.) dissolved in fresh 100 mM NaCO_3_ was applied to each chamber and left to bind for 3 h. Chambers where then rinsed thoroughly with water, dried with a N_2_ gun, and stored in an anaerobic atmosphere.

### Microfluidic flow chamber preparation

Six holes, three each side positioned to fit the chamber of the gasket, were drilled on a microscope slide (VWR) using a DREMEL 3000 rotatory tool and a 1 mm diamond coated drill tip (UKAM Industrial Superhard Tools). The previously prepared gasket coverslips were applied, aligning each chamber to the holes (one influx and one efflux). Two polypropylene tubes were allocated in each one of those (BD intradermic, efflux: 0.76 x 1.22 mm influx: 0.38 x 1.09 mm) and fixed in place with fast dry epoxy resin (Araldite). Prior to use, each chamber is filled for 30 min with a 1 mg/ml solution of Streptavidin (NEB) and then passivated with a blocking solution (20 mM Tris-HCl pH 7.5, 50 mM NaCl, 0,5 mg/ml BSA, 1% Tween).

### Single molecule imaging

All the solutions were flowed into the chamber using an infusion pump (WPI, AL-1000) with a flow rate of 30 μl/min. Before imaging, chambers were washed thoroughly with SSA (Strand Separation Assay) buffer (30 mM Hepes pH 7.6, 100 mM Potassium glutamate, 10 mM Magnesium acetate), then DNA (1 fM) was applied for 1 min. The unbound molecules were washed away with SSA buffer followed by the same solution containing oxygen scavengers (10 mg/ml Glucose oxidase, 0.4 mg/ml catalase, 1 mM Trolox, 0.8 mg/ml of D-Glucose) to stabilize the fluorophores. Images were taken using a Ti eclipse inverted microscope housing an oil immersion objective (CFI SR HP Apo TIRF 100XC Oil; Nikon). The fluorophore ATTO^647^ was excited using a 300 mW fiber laser at 647 nm wavelength (MPB communications inc) with a power intensity of 583.1 μW (±2.405 μW) at the sample and 500 ms exposure time. Fluorophore ATTO^565^ was excited using a 150 mW laser at 561 nm wavelength (Coherent) with a power intensity of 1444 μW (±0.853 μW) at the sample and 500 ms exposure time. The laser power reaching the sample was measured with a PM100 power meter (Thorlabs). The emitted fluorescence was filtered using a 405/488/561/647 nm Laser Quad Band Set (Chroma) and the signal was detected and recorder using an EMCCD iXon X3 897 camera (ANDOR technology). Images were captured every 5 seconds for 10 minutes. Protein solutions (30 μl) were diluted to a final concentration of 50 nM in SSA containing oxygen scavengers and flowed in to initiate the reaction.

### Microscopy data analysis

TIRF timelapse images were processed automatically within Fiji (Schindelin et al., 2012) for drift-correction, then maxima were assessed to define initial molecule location regions of interest. The regions were measured over time to obtain fluorescence intensity relative to the initial intensity, with drop detection triggered if fluorescence intensity drops below 75% of initial intensity for 5 sequential frames. A fully annotated ImageJ Macro is available on GitHub (https://github.com/GMerces/DisappearingSpots).

### DNA scaffolds

DNA scaffolds were prepared by adding each oligonucleotide (10 μM) to a 20 μl volume containing 30 mM HEPES-KOH (pH 8), 100 mM potassium acetate and 5 mM magnesium acetate. Mixed oligonucleotides were heated in a PCR machine to 95°C for 5 minutes and then cooled 1°C/min to 20°C before being stored at 4°C. Assembled scaffolds were diluted to 1 μM and stored at −20°C.

DNA scaffolds were imaged with a Typhoon FLA 9500 laser scanner (GE Healthcare). Cy5 labelled oligos were excited at 400 V with excitation at 635 nm and emission filter LPR (665LP). Cy3 labelled oligos were excited at 400 V with excitation at 532 nm and emission filter LPG (575LP). Images were processed using Fiji subtracting the background and adjusting the overall contrast (Schindelin *et al.*, 2012). All experiments were independently performed at least thrice and representative data is shown. The melting temperature of each probe was measured preparing a 20 μl reaction containing 0.4 μM of probe in strand separation assay buffer. A melting curve determination experiment from 20 to 95 °C was then performed on the reaction on a Rotor-GenQ qPCR machine (QIAgen) with 563 excitation filter and 610 high-pass emission filter (gain:10). The curve obtained and the melting value of each peak are reported in Table S1.

### Black Hole Quencher DNA strand separation assay (BHQ-SSA)

Each DNA scaffold containing one oligonucleotide with BHQ2, one unlabelled with Cy5 and one unlabelled (12.5 nM final concentration) was diluted in SSA buffer. Reactions were performed using a flat-bottom black polystyrene 96-well plate (Costar #CLS3694). All reactions were made in triplicate, with fluorescence output measured every 30 seconds for a total of 75 min, allowing the first point to be measured without the protein and then adding DnaA at a final concentration of 650 nM. For all reactions a negative control without protein was performed in triplicate (background). At each timepoint the average background value is subtracted from the experimental value, thus reporting the specific DnaA activity on a single substrate. All the reactions were prepared on ice to ensure the stability of the DNA probe, then allowed to equilibrate to the reaction temperature (20°C) before starting the measurement.

### Electrophoretic mobility shift assay

For electrophoretic mobility shift assays, scaffolds (13 nM) were mixed with DnaA proteins (500 nM) in 10 mM HEPES-KOH (pH 8), 100 mM potassium glutamate, 1 mM magnesium acetate, 30% glycerol, 10% DMSO and 1 mM nucleotide (ADP or ATP). Reactions (30 μl) were incubated at 20°C and after 3 hours were placed on ice for 5 minutes. 20 μl of each reaction was loaded onto a 6 % polyacrylamide (19:1) gel and run at 70 V for 6 hours in buffer containing 45 mM Tris, 45 mM boric acid and 5 mM magnesium chloride. All experiments were independently performed at least thrice and representative data is shown.

### DnaA protein purification

Plasmids encoding for His_14_-SUMO-DnaA proteins were transformed into BL21(DE3)-pLysS. Strains were grown in 2X YT medium at 37°C and at A_600_ 0.4, 1 mM IPTG was added and cultures were incubated for 4 h at 30°C. Cells were harvested by centrifugation at 7000 g for 20min, resuspendend in 40 ml of Ni^2+^ Binding Buffer (30 mM HEPES [pH 7.6], 250 mM potassium glutamate, 10 mM magnesium acetate, 30 mM imidazole) containing 1 EDTA-free protease inhibitor tablet (Roche #37378900) and flash frozen in liquid nitrogen. Cell pellet suspensions were thawed and incubated with 0.5 mg/ml lysozyme with agitation at 4°C for 1 h. Cells were disrupted by sonication (Fisherbrand 505, 60 min of 20 sec ON/OFF cycles at 30% power with a ¼ inch tip in an ice bath). To remove cell debris the lysate was centrifuged at 24,000 g for 30 min at 4°C, then passed through a 0.2 μm filter for further clarification. All the further purification steps were performed at 4°C using Fast Protein Liquid Chromatography (FPLC) with a flow rate of 1 ml/min.

The clarified lysate was applied to a 1 ml HisTrap HP column (GE), washed with 10 ml Ni^2+^ High Salt Wash Buffer (30 mM HEPES [pH 7.6], 1 M potassium glutamate, 10 mM magnesium acetate, 30 mM imidazole) and 10 ml of 10% Ni^2+^ Elution Buffer (30 mM HEPES [pH 7.6], 250 mM potassium glutamate, 10 mM magnesium acetate, 1 M imidazole). Proteins were eluted with a 10 ml linear gradient (10-100%) of Ni^2+^ Elution Buffer. Fractions containing the protein were applied to a 1 ml HiTrap Heparin HP affinity column (GE) equilibrated in H Binding Buffer (30 mM HEPES [pH 7.6], 100 mM potassium glutamate, 10 mM magnesium acetate). Proteins were eluted with a 20 ml linear gradient (20-100%) of H Elution Buffer (30 mM HEPES [pH 7.6], 1 M potassium glutamate, 10 mM magnesium acetate). Fractions containing DnaA (usually 3 ml total) were pooled and digested overnight with 10 μl of 10 mg/ml His14-Tev-SUMO protease (Frey and Gorlich, 2014).

The digested reaction product was applied to a 1 ml HisTrap HP column to capture uncleaved protein, His14-SUMO-tag and His_14_-TEV-SUMO protease. Cleaved DnaA proteins were collected in the flow-through and their purity was confirmed on SDS-page. Glycerol was added (20% final) and proteins aliquots were flash frozen in liquid nitrogen before being stored at −80°C.

### Protein crosslinking in solution

DnaA^CC^ proteins were diluted in 10 mM HEPES-KOH (pH 8), 100 mM potassium glutamate, 100 mM sodium chloride, 10 mM magnesium acetate, 25% glycerol, 0.01% Tween-20 and 2 mM nucleotide (ADP or ATP) at a final concentration of 3μM. Reactions were incubated at 37°C for 10 min, then 7.5 μl of 20 mM bismaleimidoethane (BMOE)(5.5 mM final, Thermo Fisher Scientific #803561) was added to crosslink proteins. After 6 min the reaction was quenched by addition of 15 μl of 200 mM cysteine (70 mM final) for 10 min. Samples were fixed adding 21 μl of NuPAGE LDS sample buffer (Thermo Fisher Scientific) at 98°C for 5 min and complexes were resolved and visualized by running 10 μl from each reaction following the method described below. The interpretation of the crosslinked species was based on their migration relative to molecular weight markers. All experiments were independently performed at least thrice, and representative data are shown.

### Immunoblotting

Fixed samples were resolved on a NuPAGE Novex 4–12% Tris-Acetate Gel (Thermo Fisher Scientific) then transferred to a PVDF membrane using Turbo-Blot transfer apparatus and Trans-Blot TurboTM Midi PVDF Transfer Packs (Bio-Rad). The membrane was blocked with PBS + 5% milk for 1 hour at room temperature then incubated with a PBS + 1% milk solution containing a 1:2000 polyclonal anti-DnaA antibody (Eurogentec) for 1 hour at room temperature. The membrane was washed three times with PBS + 0.05% Tween-20 and then incubated with PBS + 1% milk solution containing a 1:10.000 anti-rabbit horseradish peroxidase conjugated secondary antibody (A0545, Sigma-Aldrich). The membrane was washed three times with PBS + 0.05% Tween-20 and then incubated for 5 min with Pierce ECL Plus substrate (Thermo Scientific). Chemiluminescence was detected using an ImageQuant LAS 4000 imager. Images were processed using Fiji (Schindelin *et al.*, 2012).

### Fluorescent ATP analog binding assay

Proteins (8 μM final) were incubated with 0.16 mM of (2'-(or-3')-O-(trinitrophenyl) adenosine triphosphate (TNP-ATP) (Thermo Fisher Scientific), in 50 μl of Binding Buffer (10 mM HEPES-KOH (pH 8), 100 mM potassium glutamate, 5 mM magnesium acetate). A reaction containing 8 μM of lysozyme was used as a negative control. All reactions were performed for 10 min at 25°C in a black flat-bottom polystyrene 96-well plate (Costar #CLS3694). TNP-ATP was excited at 410 nm and fluorescence emission was detected between 500-650 nm using a plate reader (CLARIOstar Plus, BMG Labtech).

## ACKNOWLEDGEMENTS

We thank members of the Murray lab for critical review of the manuscript. We thank Frances Davison for technical assistance. Research support was provided to HM by a Wellcome Trust Senior Research Fellowship (204985/Z/16/Z).

## AUTHOR CONTRIBUTIONS

SP, DB, HY, HM conceived the research plan. SP and DB generated results presented in the manuscript. GM developed the image analysis pipeline and assisted with the image analysis. SP created Figs. SP, HY, HM wrote the manuscript. SP, HY, HM edited the manuscript.

## CONFLICT OF INTEREST

Authors declare that they do not have any conflicts of interest.

## FIGURE LEGENDS

**Figure S1.**
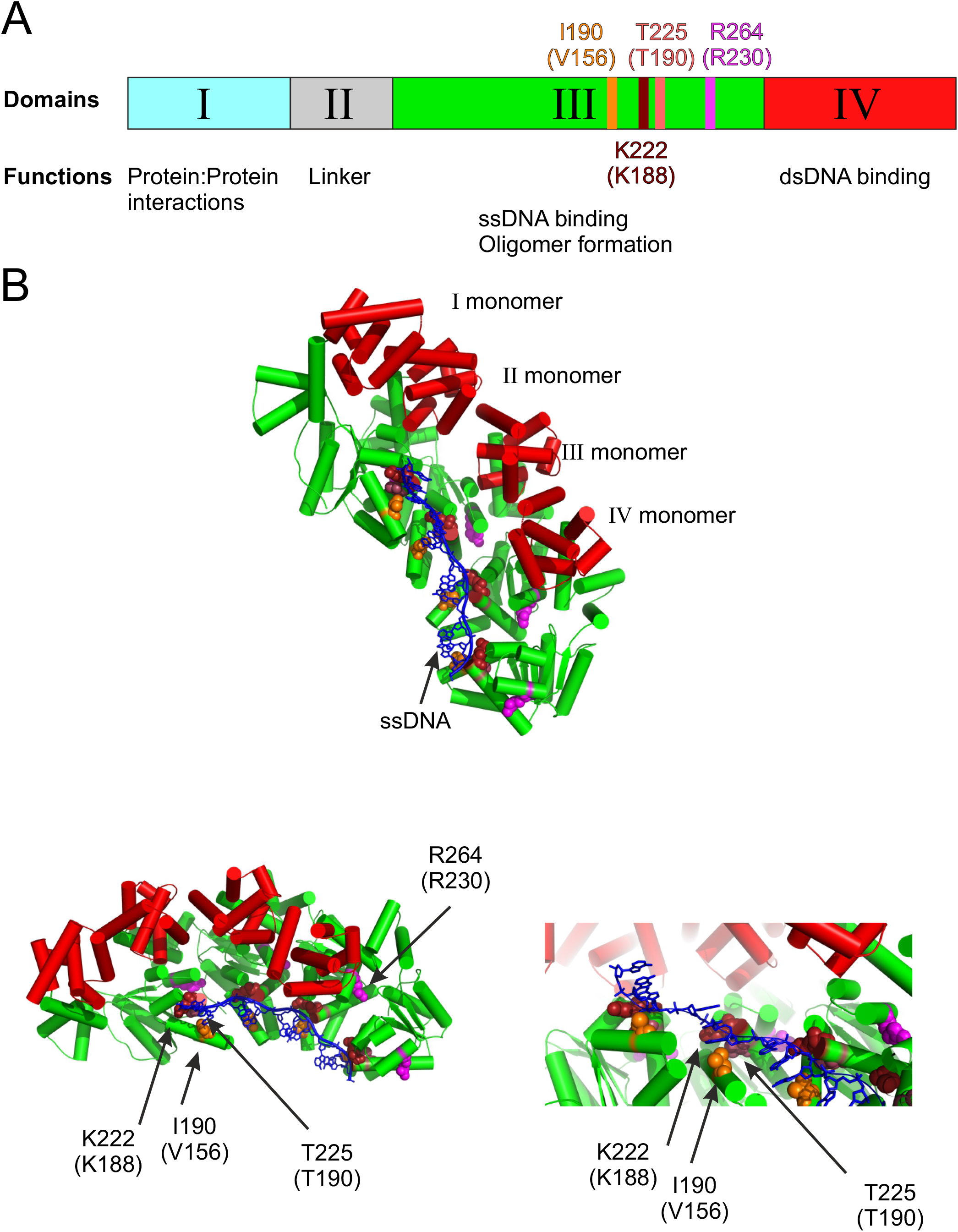
Critical residues within DnaA required for oligomerizaton and ssDNA binding. **(A)** Schematic representation of DnaA domains. Locations of critical residues are indicated. **(B)** Crystal structure of *A. aeolicus* DnaA (PDB:3R8F)(Duderstadt *et al.*, 2011) in complex with ssDNA. Locations of critical residues are indicated: arginine finger (R264), ssDNA binding (I190, K222, T225).

**Figure S2.**
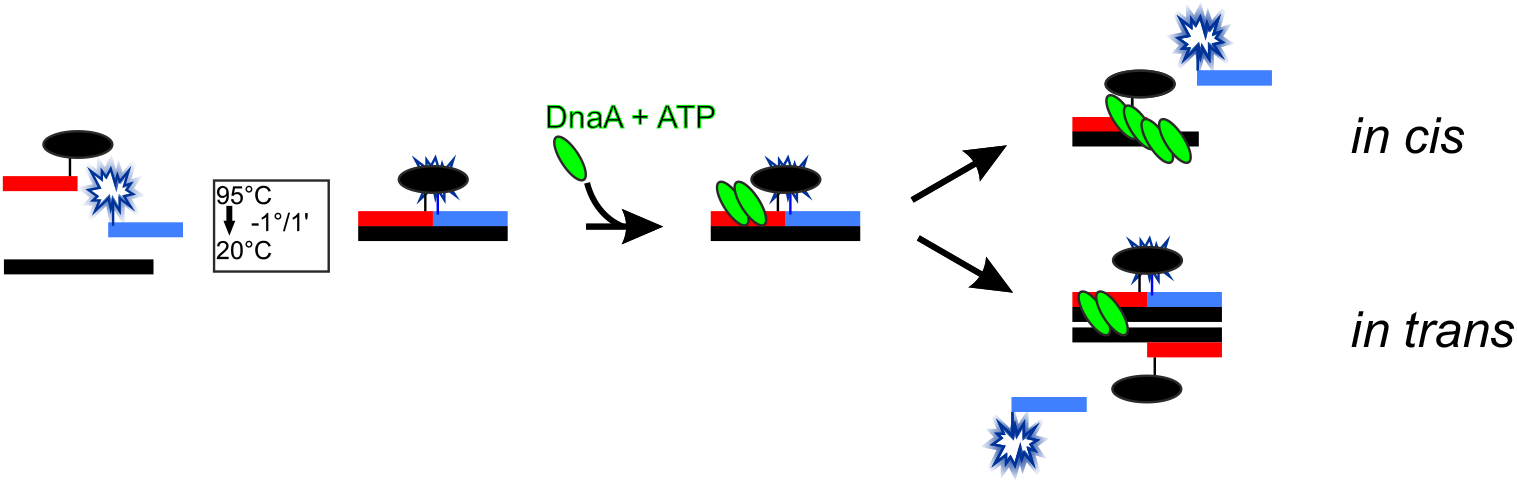
Schematic representation of the two proposed strand separation models (*in cis* and *in trans*) and their mechanism of action on BHQ-SSA.

**Figure S3.**
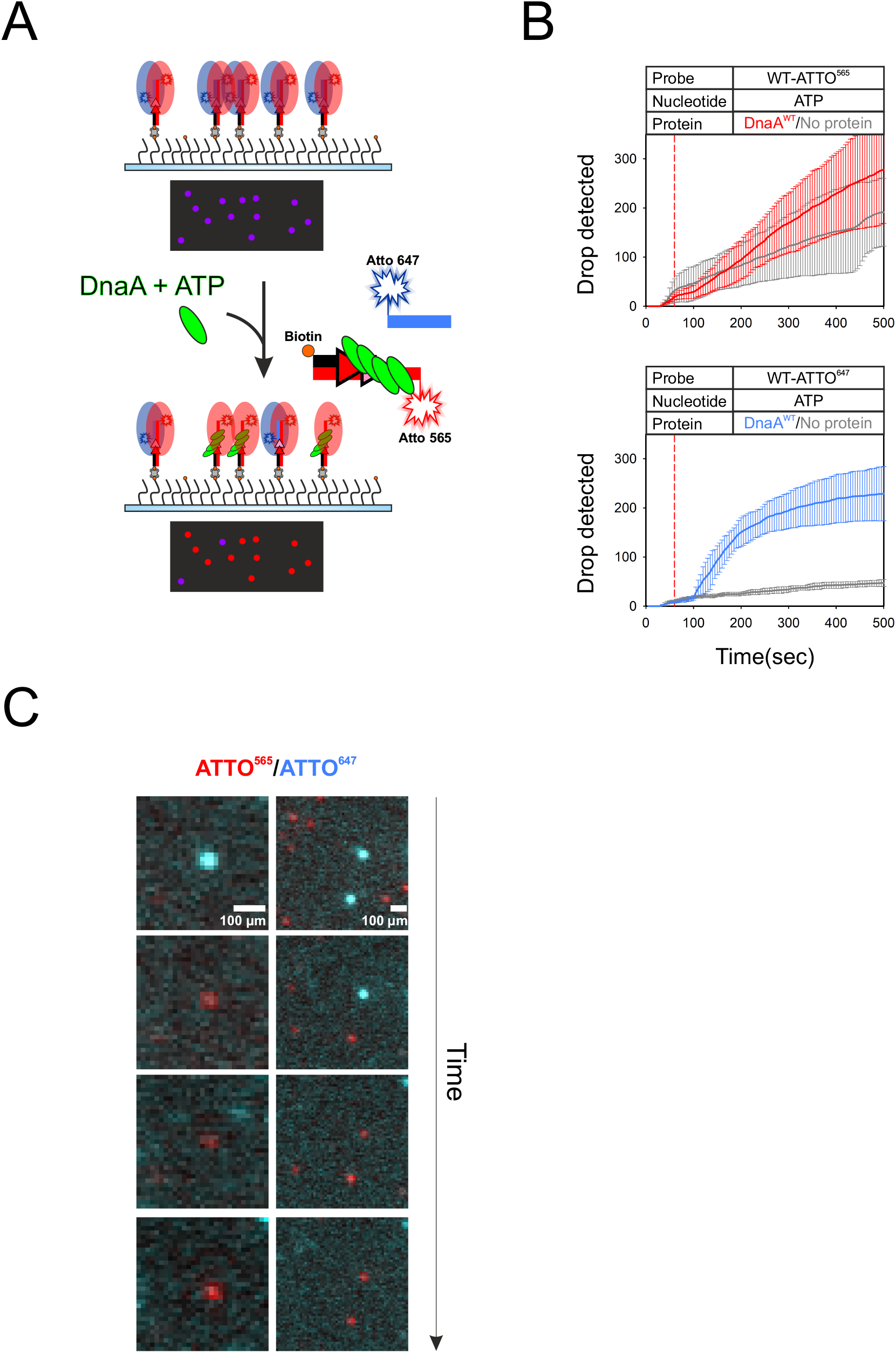
Single molecule experiments performed with an ATTO^565^/ATTO^647^ probe. **(A)** Experimental design using a DNA scaffold containing two fluorescently labelled oligonucleotides. **(B)** Assays reporting the number of colocalized spots which drop their intensity during time, observed with and without DnaA in the two channels (ATTO^565^ red, ATTO^647^ cyan). **(C)** Examples showing specific loss of ATTO^647^ fluorescence (cyan), indicating a DnaA-dependent strand separation event.

**Figure S4.**
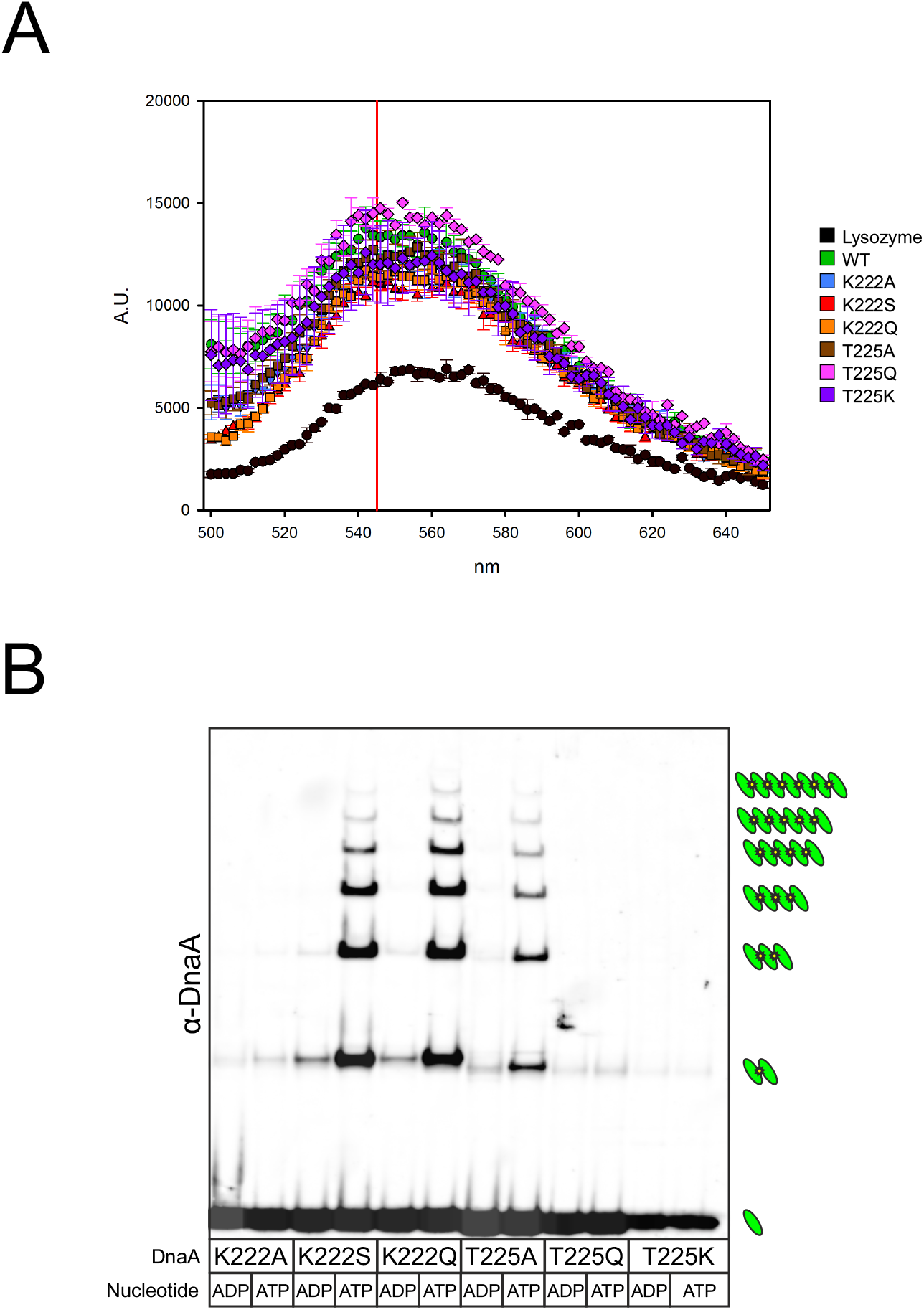
Characterization of DnaA^K222^ and DnaA^T225^ protein variants. **(A)** TNP-ATP binding assay. All DnaA proteins appear competent to bind the fluorescent ATP analog. Lysozyme was used as a negative control. The vertical red line indicates the peak emission wavelength for TNP-ATP. **(B)** Filament formation assay in solution performed with the several DnaA proteins. Crosslinked species are indicated to the right.

**Figure S5.**
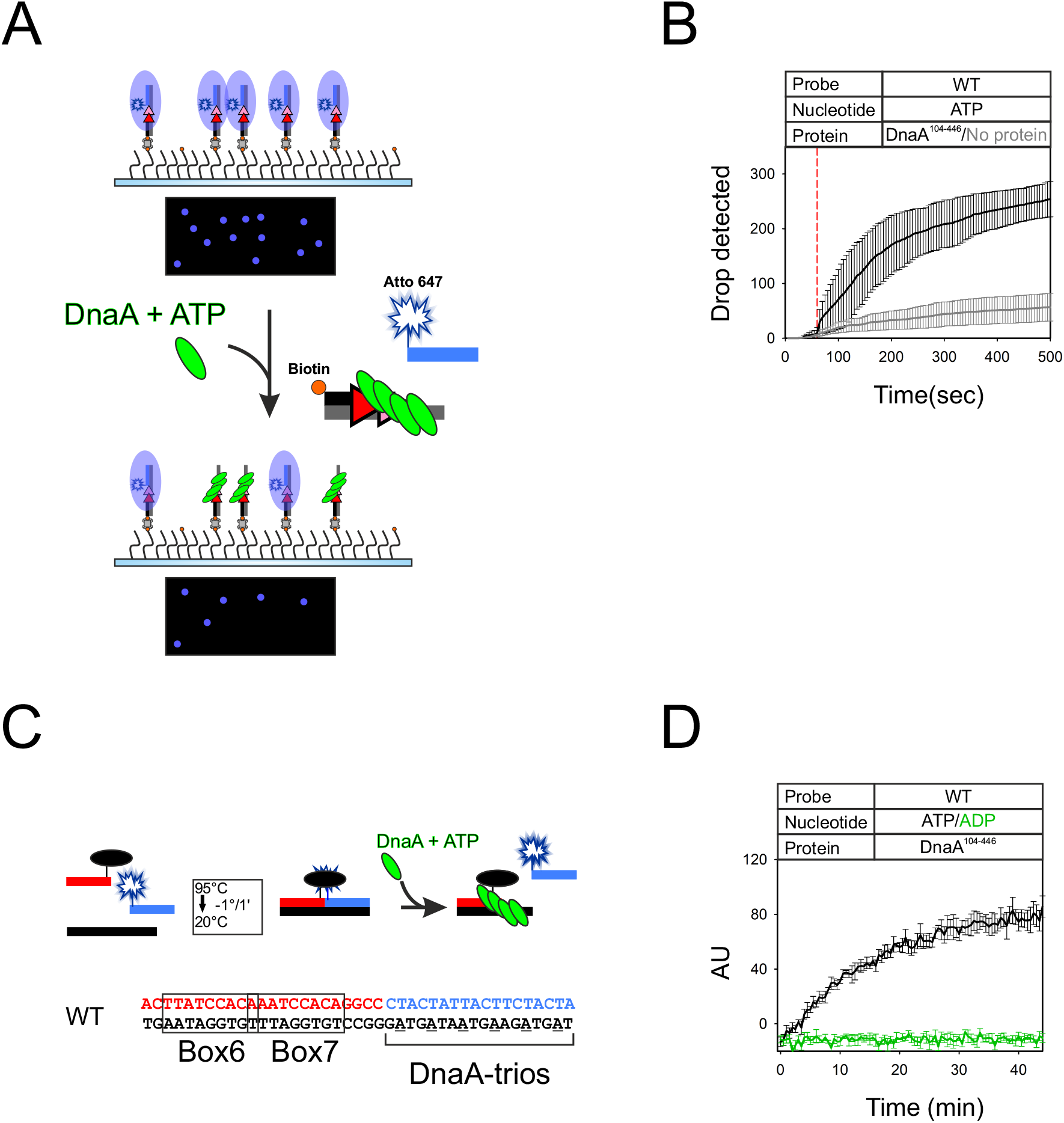
DnaA domains III-IV and necessary and sufficient for DNA strand separation activity. **(A)** Experimental setup of single molecule strand separation assay performed using a DnaA truncation lacking domains I-II (DnaA^104-446^). **(B)** Graph representing the additive number of spots with a fluorescent signal drop over time is reported on the right. **(C)** Schematic representation of BHQ strand separation assay. The wild-type BUS sequence is shown for comparison. **(D)** Graph showing the assay performed with DnaA^104-446^ with ATP or ADP and the relative increase in fluorescence during time.

**Figure S6.**
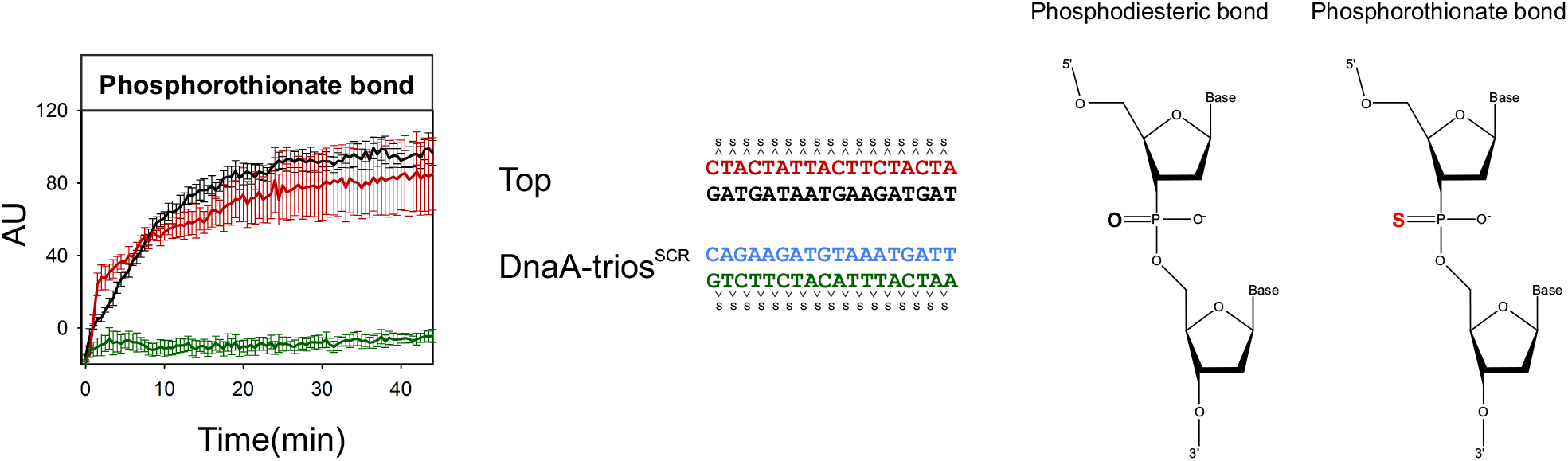
Phosphorothionate bond impairment is dependent by their presence on the DnaA-trios backbone. BHQ-SSA performed with substrates harbouring phosphorothionate modifications on the displaced strand backbone or on the DnaA-trios^SCR^ one. Positions of chemical modifications are indicated to the right.

## Notes

### Competing Interest Statement

The authors have declared no competing interest.

